# On the potential sources of a low-frequency sound percept only a few can perceive

**DOI:** 10.1101/2025.06.07.658448

**Authors:** Bonifaz Baumann, Andrej Voss, Carlos Jurado, Markus Drexl

## Abstract

A small percentage of the general population reports almost constant humming or pulsing low-frequency sound percepts (LFSPs) while others in their vicinity, such as family members, often do not perceive these sounds. The origin of these LFSPs remains to be elucidated and may or may not be related to external sound sources. The underlying causes of these perceptions could also be subjective and belong to the tinnitus family, especially in cases where no external sound source sufficiently explaining the LFSP can be found. The present study puts forth two hypotheses to explain the phenomenon, based on both subjective and objective auditory phenomena: an unusually high auditory sensitivity to low-frequency sound, and hearing one’s own low-frequency spontaneous otoacoustic emissions (SOAEs), sounds actively produced by the inner ear as a normal, physiological by-product of cochlear amplification. The present study employed high-resolution, low-frequency hearing threshold measurements and SOAE measurements in 28 individuals with LFSPs, and in control groups devoid of LFSPs. LFSP complainants self-reported hearing a LFSP at a median frequency of 50 Hz obtained with a frequency-matching procedure. With a few clear exceptions, complainants most often did not present unusually sensitive low-frequency hearing thresholds. Furthermore, hearing threshold fine structure was comparable to the control group. In addition, no SOAEs in the low-frequency range could be measured. Based on our results, while cases of hearing physical external sound sources are not ruled out, we suggest that a subjective tinnitus in the low-frequency range is often the reason behind hearing these LFSPs.

## Introduction

A small percentage of the general population, estimated to be between 2% and 4%, report an almost constant perception of a low-frequency sound. The acoustic quality of this auditory phenomenon is often compared to the sound of an idling diesel engine [1]. The characteristics of these low frequency sound perceptions (LFSPs), frequently referred to as the “hum phenomenon”, vary considerably between individuals and can be described as pulsing, humming, rumbling, or throbbing [1, 2]. A distinctive attribute of the hum is its tendency to be perceived exclusively by specific individuals, while others within the same familial unit or geographical vicinity often may not be able to hear it. The hum phenomenon has in the past years gained attention due to an increase in reports of LFSPs in the daily press in specific geographical regions, such as Oslo, Norway [3], Darmstadt, Germany [4], or Windsor, Canada [5], and others. LFSPs have been reported on various continents, and a map collecting the geographical distribution and characteristics of reported LFSPs is available on a dedicated website [6]. It is important to consider the possibility of reporting bias when drawing conclusions from the reported cases, which appear to show clear geographical clusters. Several studies have attempted to identify external sound sources as the cause of LFSP. However, potential external sound sources (ESS) were only reported in a few studies [7, 8]. Despite the absence of a definitive explanation for this phenomenon, three attempts to elucidate its relation to the auditory system are reviewed briefly below:

### The perception of external low-frequency sounds in subjects with unusually sensitive low-frequency hearing thresholds

LFSPs were reported in both the low-frequency and infrasound ranges, but typically with frequencies ranging from about 30 to 80 Hz [1]. Low-frequency sound (LFS) is loosely defined as sound with frequencies below 100-200 Hz [9, 10], while infrasound is typically defined as sound with frequencies below 16-20 Hz [9, 10], a frequency range which was originally considered to be inaudible to humans. However, several studies have shown that these frequencies can be heard when sound levels are sufficiently high, and that there is no sudden perceptual transition between LFS and infrasound [10]. Nevertheless, *normal* low-frequency and infrasound hearing thresholds (LFHTs) are much higher than those in the normal human hearing frequency range [10]. Most people seem to not experience annoying LFSPs, despite abundant low-frequency noise pollution in the industrialised world [e.g. 11]. However, a small percentage of the general population appears to have unusually low LFHTs, and considerable individual differences in hearing sensitivity, especially in the low-frequency range, exist [12], a result of the estimated normal distribution of normal hearing thresholds. At 31.5 Hz, for example, there is an approximately 17-dB difference between the first percentile and the median of hearing thresholds from a normal-hearing cohort [13]. As an example, for a place like Trondheim, Norway with its ca. 250,000 inhabitants, this means that 2500 people will hear a 31.5 Hz sound with a sound pressure level of 43 dB SPL (ca. 17 dB lower than the ISO389-7:2019 threshold for that frequency), and 247,500 people will not. Consequently, it can be speculated that low-frequency sound hearers (LFSHs), a term we use here to refer to subjects hearing a LFSP, have hearing thresholds belonging to at least the lower half of the distribution of normal hearing thresholds. It has long been known that normal hearing thresholds show also a prominent microstructure, characterised by considerable threshold changes (up to 20 dB) associated with very small changes of test tone frequency [14, review: 15]. Such local hearing threshold maxima and minima go often unnoticed during routine clinical hearing tests due to the limited number of frequencies tested and the large spacing between, and threshold microstructures might represent another explanation for LFSPs [16, 17]. While other sensory systems such as the vestibular system and the somatosensory system can also be stimulated by sound, sound does not represent the adequate stimulus for these sensory systems, and high stimulus intensities are required to exceed detection thresholds [10, 18–20]. In addition, to the best of our knowledge, sensory systems other than the auditory system cannot mediate the sensation of pitch, but most LFSHs seem to be able to characterise the pitch of their LFSP at least to some extent [17]. It is therefore quite unlikely that other sensory systems contribute to the perception of LFS, except for sound presented at high intensities.

### Spontaneous otoacoustic emissions (SOAEs), sounds actively produced by the inner ear, which are heard by their owners

SOAEs are sounds produced by the inner ear as a by-product of physiological processes [see 21 for review]. The exact mechanisms behind the production of SOAEs are still unclear, but two prevailing theories regarding their origin exist, both involving active outer hair cell (OHC) motility in the Organ of Corti of the inner ear [see 22 for review]. It has been estimated that approximately half of the population exhibits measurable SOAEs, with a trend of higher prevalence among female subjects [21, 23, 24]. SOAE frequencies are typically situated between around 0.5 and 6 kHz [21] and, to the best of our knowledge, low-frequency SOAEs have not been reported in humans. SOAE sound levels measured in the closed ear canal typically range from about −10 to about 20 dB SPL [21]. Consequently, a considerable proportion of SOAE levels may exceed the hearing threshold of their owners. This assertion is particularly valid considering the expectation that the intracochlear sound energy is greater than that measured in the ear canal as SOAE sound pressure levels [25]. SOAEs demonstrate often notable stability in terms of both frequency and level over a considerable part of their owners’ lives, as evidenced by longitudinal studies [26, 27]. However, SOAES are mostly not perceptible to their owners [23, 28], as the auditory system appears to be able to adapt to, and filter out, these persistent sounds [25, 29], or the sound pressure levels involved do simply not exceed the hearing threshold. However, SOAEs may be heard temporarily when transient changes in their frequency or level occur [25]. Reports of annoying, long-term perception of, albeit not low-frequency, SOAEs can be found in the literature [29]. Consequently, the experience of perceiving one’s own low-frequency SOAE could offer an alternative explanation for the percepts LFSHs hear, which could be regarded as objective tinnitus (see below).

### Low-frequency tinnitus

Tinnitus is a generic term that includes objective tinnitus, i.e. the perception of sounds generated within the human body, and subjective tinnitus, which is the perception of a sound in the absence of a physical sound source [30, 31]. Possible explanations for objective tinnitus include myoclonus of the middle ear muscles (i.e. brief involuntary twitching or jerking of the muscles of the middle ear or Eustachian tube), and vascular or tube malformations. These conditions are typically described by sound characteristics that are quite different from typical LFSP reports, such as being synchronous with breathing or heartbeat, or, in the case of myoclonus, manifesting as a clicking or fluttering sound [31, 32]. The underlying pathophysiology of subjective tinnitus, however, is far from being fully understood. The prevailing view is that it involves intricate interactions at all levels of the auditory system, including the cochlea, auditory nerve, and auditory cortex, as well as non-auditory systems [33, 34]. The onset of tinnitus is often preceded by peripheral, cochlear damage, but this is not necessarily the case. Tinnitus can also be triggered by stress and is influenced by the somatosensory system [34, 35]. Tinnitus is commonly thought of as a continuous, high-pitched sound, but a variety of spectral and temporal shapes have been described [36]. Those experiencing typical subjective tinnitus localise the sounds they hear mostly to the inside of their head, they *internalise* the sound [36], indicating that the sounds they perceive are not realistic. Some evidence in the literature reporting cases of tinnitus matched with frequencies as low as 29 Hz can be found [36, 37], and LFSPs and tinnitus are thus not mutually exclusive.

### Scope of this study

The present study is concerned with the origins of LFSPs, employing widely accepted biomedical methodologies. Hypotheses that do not originate in the auditory system were not evaluated or taken into consideration in this study. Consequently, the term LFSP is employed in lieu of “the hum phenomenon” due to the absence of a definitive and well-defined definition. In this context, LFSP signifies the perception of low-frequency sounds that are not audible to most individuals, irrespective of the presence of a discernible physical sound source.

## Methods

A total of 28 LFSHs were included in the study, with 13 males, 14 females, and one LFSH identifying as gender diverse. The median age of the participants was found to be 53.5 years. These participants were recruited from a social media group, where LFSHs self-organised. Additionally, 38 young, normal-hearing adults, recruited from a poll of the department’s students, without any dependencies related to the authors of this study, were divided into two control groups (CG1 and CG2).

The study was approved by the local Ethics Committee of the University Hospital Munich, Ludwig-Maximilians-Universität München, Germany (project number 19-107), in agreement with the latest revision of the Code of Ethics of the World Medical Association (Declaration of Helsinki). All participants were provided with written information about the study and its procedures and gave their written informed consent. The recruitment period lasted from 1^st^ of April 2019 until 31^st^ of May 2019.

Control group participants did not report hearing LFSPs and had no self-reported history of neuro-otological diseases. CG1 consisted of 31 participants (23 females, 8 males, median age 24 years, whereas CG2 included 7 participants (3 females, 4 males, median age 22 years). All participants were seated in a recliner placed in a sound-attenuated chamber while taking part in a set of experiments, which took in total around three hours. Breaks were given as needed. All participants underwent conventional air-conducted, pure-tone audiometry to assess their hearing thresholds in the frequency range between 125 Hz and 8 kHz before the actual experiments. The controls groups were comprised exclusively of individuals with normal hearing, defined as hearing thresholds at each tested frequency that were at least 20 dB HL, or better, but LFSHs included in the study regardless their hearing status. Conventional audiometry was conducted using commercial software (Automatic Pure Tone Audiometry V2.28, Hörtech, Oldenburg, Germany) and a gamepad (Bigben Interactive, Lesquin, France) as a feedback device, testing both ears in all participants. Test tones were presented using circumaural headphones (HDA 200, Sennheiser Electronic GmbH & Co. KG, Wedemark, Germany), and an external soundcard (Fireface UC, RME, Haimhausen, Germany) connected to a laptop (ASUS, G60VX, Taipei, Taiwan) was employed for D/A conversion, set to a sampling rate of 44.1 kHz and a word length of 24 bits. The sound system was calibrated using the software’s built-in calibration routine, employing an ear simulator (Type 4153, Brüel & Kjær, Nærum, Denmark) connected to a calibrated measuring amplifier (type 2636, Brüel & Kjær, Nærum, Denmark), to ensure correct sound pressure level over the frequency range used.

### Questionnaire and self-testing of low-frequency sound percept frequency

Prior to the measurements in our laboratory facilities, LFSHs were asked to complete a questionnaire. The questionnaire consisted of 22 questions, including numerical analogue scales, multiple choice questions, and individual response questions. The questionnaire was designed to collect demographic data about the participants, the characteristics of the LFSP, and the factors influencing it. LFSHs were also instructed to determine their sound percept frequency (SPF) at home by undertaking a frequency matching procedure using a freely available online pure tone generator where frequencies could be adjusted in 1_Hz steps. The participants were asked to adjust the frequency of a pure tone stimulus so that it would best match their SPF. They were also encouraged to use circumaural headphones if possible. LFSHs were informed of the possibility of binaural beats (that might occur when the generated tone frequency is close to their SPF), which is a spatial hearing sensation that can occur when listening to a dichotic signal that differs slightly in frequency between the ears, and they were asked to repeat the SPF matching process several times. Whilst this procedure is undoubtedly less controlled than those conducted in the laboratory, it avoids the problem that LFSPs can wax and wane and therefore may not be perceived while participants are in the laboratory facilities, rendering a frequency-matching procedure impossible. Estimated SPFs below 15 Hz were considered unlikely to be accurate due to potential equipment distortions. This is because considerable high sound pressure levels (SPLs) are required for the stimuli to be audible at such low frequencies. These are difficult to reproduce without significant distortions with consumer-grade equipment. All participants were asked if they could perceive the hum while they were in our laboratory facilities (for the most, located away from their hometown), which was confirmed by 11 participants (39% of all LFSHs).

### Measuring low-frequency hearing thresholds

#### Average LFHT from a normal-hearing cohort and high-frequency resolution LFHTs

High-frequency resolution LFHTs were measured with the intention to assess low-frequency hearing and threshold microstructure in the LFSH group with pure tones in a 16-Hz range symmetrically distributed around the SPF as a centre frequency. LFHT measurements were carried out at nine individual pure tone frequencies corresponding to the SPF, SPF ±1 Hz, SPF±2 Hz, SPF±4 Hz, and SPF±8 Hz, respectively. If participants were unable to match their SPF or if matching it required frequencies below 15 Hz, thresholds were measured at seven default frequencies of 16, 20, 40, 63, 70, 80 and 125 Hz, which means that threshold microstructure could not be assessed. Threshold measurements were carried out in one ear only. If the participants indicated that the LFSP could be lateralised to one ear, that ear was measured, and if not, one ear was randomly chosen. 14 left ears and 14 right ears were measured. The same seven default frequencies were also used for threshold measurements in CG1 participants (from one randomly selected ear only (20 right ears and 11 left ears). The data from CG1 was used to develop an average LFHT from a normal-hearing cohort, not reporting LFSPs. To be able to compare the LFSH thresholds obtained with individually varying test tone frequencies (which depended on the self-reported SPF) to the normal-hearing average LFHTs (obtained with default test tone frequencies), a threshold value for each frequency point used in the LFSH group measurements was required. To achieve this, the CG1 hearing thresholds were fitted with a polynomial function (3^rd^ order, red line in Fig 2; MatLab’s “polyfit” function was used) to obtain an extra- and interpolated LFHT. A reference range (RR) of normal LFHTs was obtained by calculating the 2.5^th^ and 97.5^th^ percentile of the differences between the estimated, mean hearing thresholds (from interpolation) and the measured hearing thresholds, at each measured frequency. The upper limit (97.5^th^ percentile) of this RR was accordingly calculated to be 11 dB above the fitted LFHT, while the lower limit (2.5^th^ percentile) was 12.2 dB below, resulting in a total RR of 23.2 dB around the fitted LFHT. In addition, high-frequency resolution LFHTs were also measured in seven participants of CG2 with randomly selected, individual sets of frequencies used for participants in the LFSHs group. Those participants were measured several times and contributed 20 high-frequency resolution LFHTs in total.

#### Technical setup and stimulation

All stimuli were time-windowed with cos^2^ ramps to avoid onset-offset transients, and the stimulation parameters were adjusted as a function of frequency. Stimulus parameters were adapted from Kuehler, Fedtke (38) and are summarised in Table 1. Since LFSHs were tested at individual frequencies, depending on their SPF, stimulus parameters were interpolated or extrapolated where needed, based on the parameters shown in Table 1.

**Table 1.**
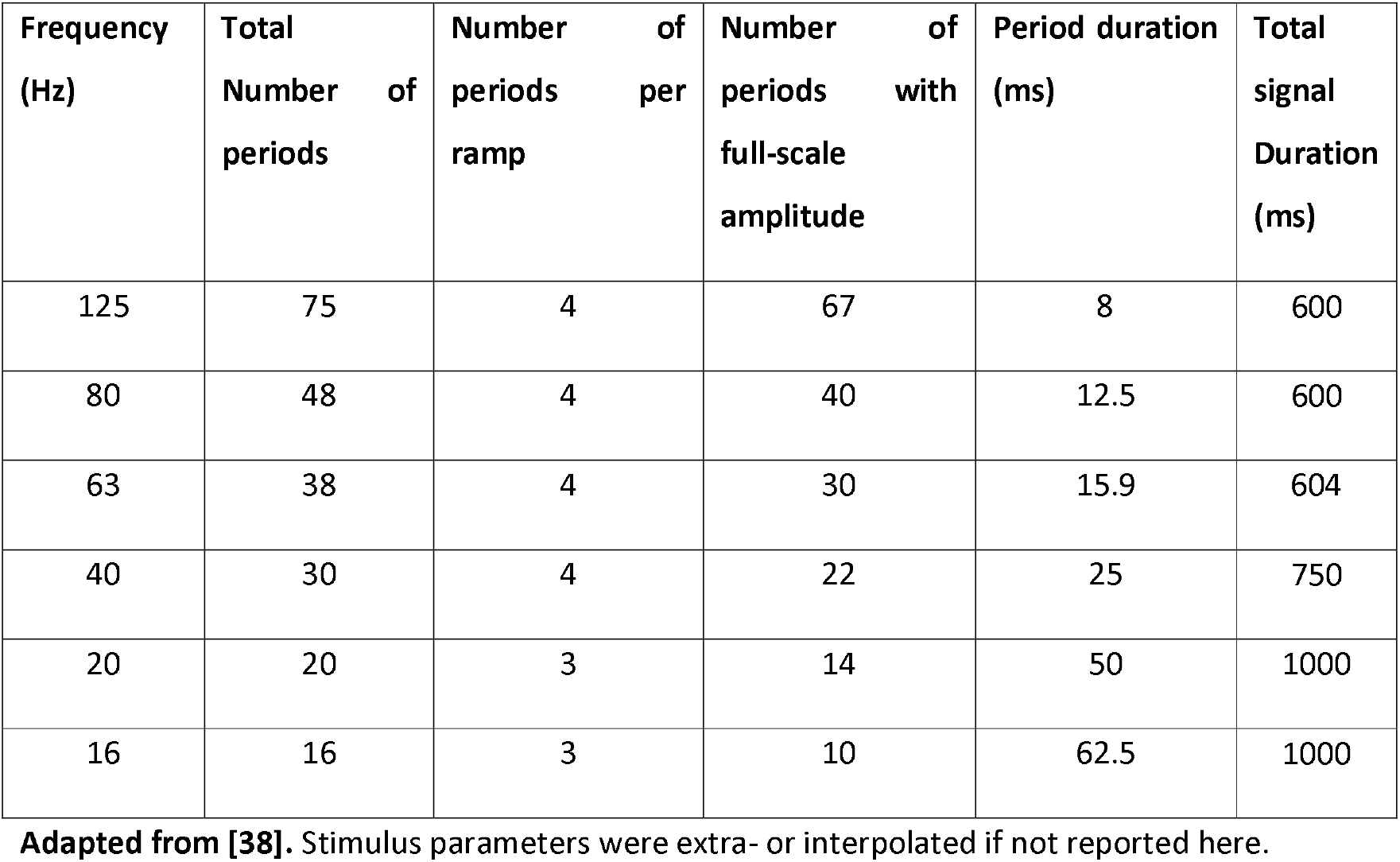
Stimulus parameters for measurements of low-frequency hearing thresholds.

The experimental procedures were controlled, and the signals used were generated using scripts custom-written in MATLAB (RRID:SCR_001622) 2015a (The MathWorks, Inc., Natick, Massachusetts, USA) with the SoundMexPro toolbox [39]. Experiments were run on a laptop connected to an external soundcard, as mentioned above). The generated analogue signals, required also for calibration, were amplified using an AV receiver (RB-971, Rotel, Tokyo, Japan) and sent to a modified aluminium diaphragm loudspeaker (Rockwood 8” subwoofer, Sintron GmbH, Iffezheim, Germany) placed outside the sound-attenuated chamber. An acrylic glass plate with a 3-cm opening in the centre covered the loudspeaker. The latter was acoustically connected to the ear of the participant (who sat inside the sound attenuated chamber) via an 8.8-m long tube, sealed at one end into the loudspeaker’s opening and going through an opening in the chamber’s wall. The tube diameter gradually decreased and at its other end went through a pierced foam earplug (E-A-R Classic II PD01200, 3M, Nadarzyn, Poland), thereby delivering the LFS stimuli into the ear canal. The earplug had an additional perforation where another tube was inserted to connect a microphone (ER-10C, Etymotic Research Inc., Elk Grove Village, IL), used for in-situ calibrations in the participants’ ear canal. To ensure accurate sound levels, the sound system was also calibrated beforehand using an ear simulator (Type 4157, Brüel & Kjær, Nærum, Denmark) and a calibrated measuring amplifier (Type 2636, Brüel & Kjær, Nærum, Denmark).

#### Psychoacoustic procedure

The LFHT measurement paradigm consisted of a one-up one-down staircase procedure used in combination with an adaptive two-alternative forced-choice paradigm (modified from [38]). Participants used a touchscreen, connected to a laptop situated outside the soundproof chamber, as a feedback device. On the touchscreen, two buttons were sequentially highlighted and represented two intervals. One of these intervals was randomly accompanied by the test stimulus, whereas the other was accompanied by silence. Participants were tasked with the identification of the interval in which the LFS was present, by pressing the corresponding button. The frequency-dependent stimulus start levels were adopted from [38], and for stimulus frequencies not described there, data interpolation was used. Based on the participants’ response, the sound pressure level was adjusted accordingly: when the participants responded correctly or wrongly, the stimulus level was lowered or increased, accordingly. This process created reversal points, indicating transitions from wrong to correct responses, and vice versa. The measurements were concluded after completing 12 reversal points, and the LFHT was estimated by averaging the sound pressure levels corresponding to the last eight reversal points. To ensure precise adjustments, stimulus level increments and decrements were gradually reduced. At the beginning of the measurement, level increments were set to 12 dB, and level decrements to 4 dB. Level steps decreased progressively with increasing number of reversal points until a step size of 3 dB for increments, and 1 dB for decrements was reached. Before reaching the first reversal point, the participants could indicate that the initial LFS presentation level was below their LFHT, if needed, prompting an increase in start level by 4 dB. This option was no longer accessible once the first reversal point was reached. If the maximum amplitude difference between the last eight reversal points exceeded 15 dB, indicating inconsistency in the participant’s responses, the data was not included for analysis, and the measurement was repeated. This was necessary for at least one measurement in 23 of the 28 LFSHs, with a total of 33 repeated measurements (out of 234 measurements in total).

### SOAE testing

The experimental setup used for recording spontaneous otoacoustic emissions (SOAEs) in this study was identical to that used in a previous study (Kugler et al., 2014). However, in the present study, two ER-10C DPOAE probe systems (Etymotic Research Inc., Elk Grove Village, IL, USA) were simultaneously inserted into both ear canals. This approach reduced the required recording time and allowed us to remove SOAE candidates with similar frequencies, recorded in both ears, which were then considered to be artefacts. SOAE candidates recorded in both ears were excluded from the analysis if they were similar within a 10 Hz range, as it is improbable (although possible, see [40]) that SOAE would occur at very similar, or identical, frequencies in both ears of a participant. Signal generation - used here only for assessing the probe fit in the ear canal- and data acquisition were controlled with the same software and hardware as described in section 2. The microphone signals from the ER-10C probes were amplified by 30 dB using the microphone preamplifier of the external sound card. To maximise the detection of SOAEs, each measurement began with a 30-second trial to ensure favourable measurement conditions, excluded from the subsequent analysis. This was followed by a 2-minute measurement, which was included in the analysis. In cases where a high level of background noise occurred (mostly due to participants movement during the recording), the measurements were repeated, as an automatic evaluation of the signal-to-noise ratio (SNR) would reject SOAEs candidates in the subsequent analysis (see details below). Automatic identification of SOAEs followed the approach explained in detail in previous publications from this laboratory, e.g. [41], and the most essential steps are summarised here:

The magnitude spectrum of each recording was calculated using MATLAB’s pwelch function, implemented with a 0.5 s Hann window, 12.5% overlap, and a zero-padded FFT with a size of 1 s, resulting in a frequency bin width of 1 Hz. Candidate SOAEs were automatically detected in the magnitude spectrum by finding local maxima (minimum level: −15 dB SPL; minimum spectral distance between two maxima: 10 Hz) within a frequency range between 30 Hz and 5 kHz. SOAEs were considered valid if the power of the dominant SOAE spectral line differed significantly from the power of the noise floor (F-test, critical value: 5.39; see Dobie and Wilson (1996)). The noise floor power was calculated as the averaged power in two 8-Hz wide frequency bands, situated 20 Hz above and 20 Hz below the peak frequency of the candidate SOAE. This produced a list of SOAEs from both ears for each participant.

As the noise floor is generally inversely proportional to frequency, it is expected that low-frequency SOAEs will be challenging to measure without attempts to improve the SNR. To achieve this, an additional measurement and analysis technique was used in all participants. A dual-microphone OAE probe (Path medical GmbH, Germering, Germany) was used, and signal acquisition followed the same procedure as described above for the two microphone signals. To improve the SNR, the cross power spectral density [42] of the two recorded signals from the same ear was calculated using MATLAB’s cpsd function, and SOAE candidates were identified in the resulting signal using the same procedures as described above. While this reduces electrical, non-coherent noise originating from the two recoding pathways resulting in improved SNRs, the methods would not reduce coherent acoustic noise picked up by the two microphones. Newly detected SOAEs were only added to the list of SOAEs if their frequency differed by more than 10 Hz from already identified SOAEs (previously obtained using the two ER-10C probes). Measurements with the dual-microphone probe were carried out unilaterally in the corresponding ear of individuals who reported unilateral LFSPs, while both ears were tested on those not lateralising their LFSP, and in all tested CG2 participants.

### Statistical analysis

The statistics and machine learning toolbox was used in MATLAB 2024b (The MathWorks, Inc., Natick, Massachusetts, USA) for statistical analysis. The research questions are based on a between-subjects design (differences between the LFSH group, and the control groups). The threshold microstructure depths (dependent variable) obtained with different test tone frequencies (independent variable) was compared between the LFSH group and a control group by using a two-sided Wilcoxon rank sum test, employing the “ranksum” function. Furthermore, the distribution of microstructure minima and maxima (dependent variables), obtained as a function of test tone frequency relative to the SPF (independent variable) was compared between the LFSH group and the control group with a two-sample Kolmogorov-Smirnov test, using the “kstest2” function. The threshold for significance was α=0.025 (adjusted for multiple comparisons according to the Bonferroni correction).

## Results

### Audiometric results, participant-operated frequency matching of the low-frequency sound percept, and questionnaire results

#### Audiometric results

11 LFSHs (39.3%) had conventional audiometry hearing thresholds of 20 dB HL or worse at one or more test frequencies between 125 Hz and 8000 Hz in at least one ear, which is an expected finding given the age range of the group tested. All control group participants had hearing thresholds better than 20 dB HL at all tested frequencies.

#### Participant-operated frequency matching

Across all the individuals where it could be matched (N=23), the median SPF was 50 Hz (see Fig 1 for the distribution of self-reported LFSP frequencies). Five participants were unable to match their SPF, stating that its frequency composition was too complex to be matched with a pure tone. This number includes one participant who reported an LFSP frequency of 10 Hz, which was deemed unrealistic (see methods). Approximately 45% of the participants reported that they had attempted to measure a potential physical source of the LFSP, either themselves or with professional assistance. Approximately 20% of the participants self-reported that they were able to confirm such a physical source.

**Fig 1.**
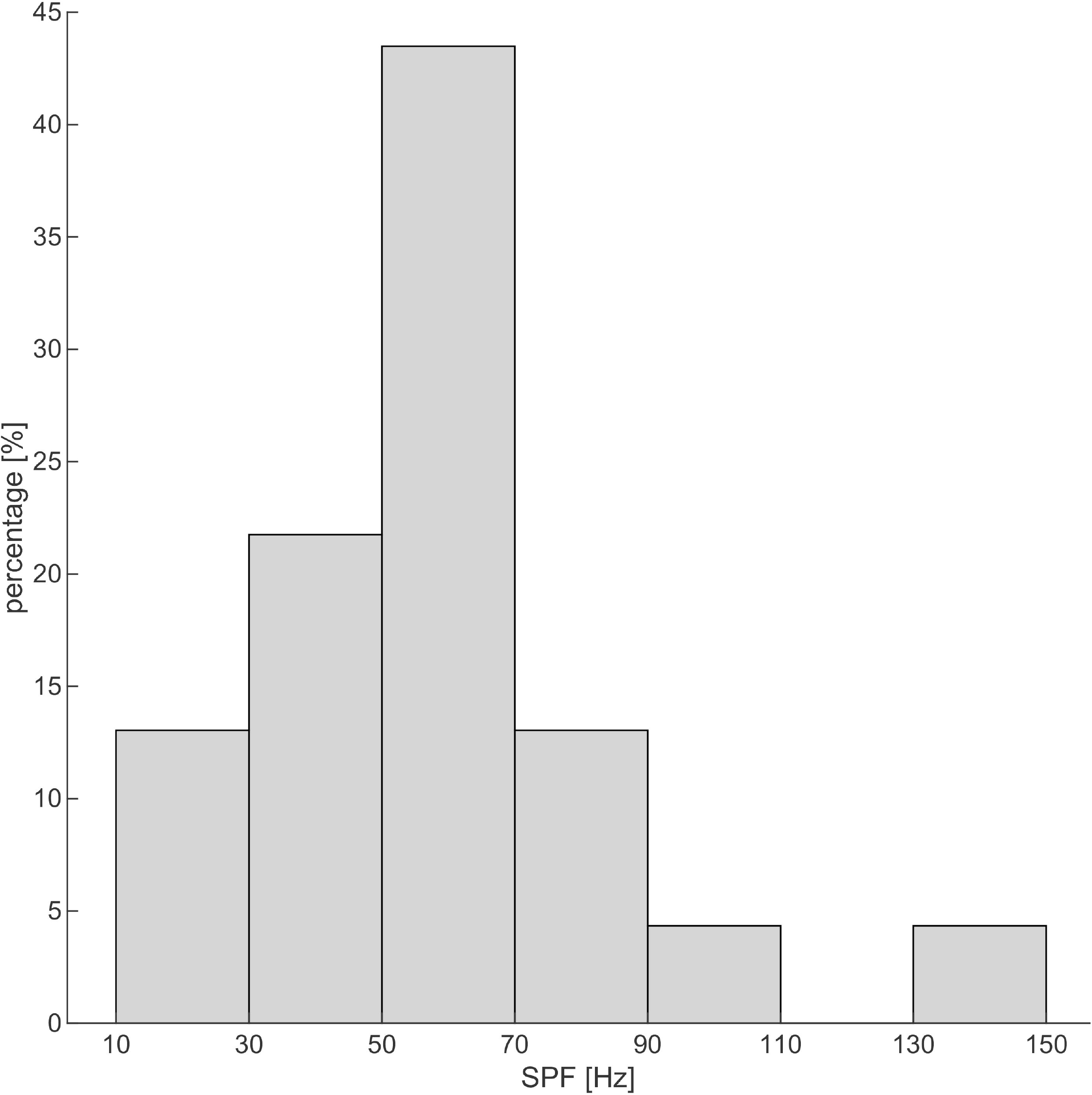
Distribution of the self-reported SPFs from 23 participants (bin width 20 Hz). The median SPF was 50 Hz.

#### Questionnaire results

68% of the LFSHs reported that they cannot lateralize the LFSP. The remaining participants were almost evenly divided into two groups, with four individuals reporting lateralized LFSPs to the left ear and five individuals indicating LFSPs in the right ear. 86% of the LFSHs classified the LFSP as stressful. Additionally, 68% of the participants responded that family or household members were unable to hear the LFSP. 39% of the participants reported hearing additional sounds of unknown origin, described by them mostly as high-frequency tinnitus. One participant had previously experienced high-frequency tinnitus in conjunction with an episode of sudden hearing loss. Most participants rated their typical acoustic environment as rather quiet (71% chose a value lower than 5 on a linear scale from 1 (very quiet) to 10 (very noisy)), with a mean score of 4.2. 68% of the participants reported that the LFSP was only audible in specific locations, and for 85% of the participants, it was not dependent on the time of day. Two participants (7%) reported that the percept was dependent on both time and place. For the four participants who described that the LFSP depended on the time of the day, the phenomenon was predominantly perceivable during the mornings and evenings. Most of the participants (ca. 93%) did not experience pitch changes of their LFSP, whereas ca. 60% described the perceived loudness of their LFSP as constant since their first encounter. The LFSP was not a constant phenomenon for ca. 45% of the LFSHs in this study, and 36% of the LFSHs indicated that the number of LFSP episodes was increasing over time. Participants were also asked to respond to a series of factors in relation to their ability to enhance or attenuate the LFSP (several answers could be given, see Fig 2).

**Fig 2.**
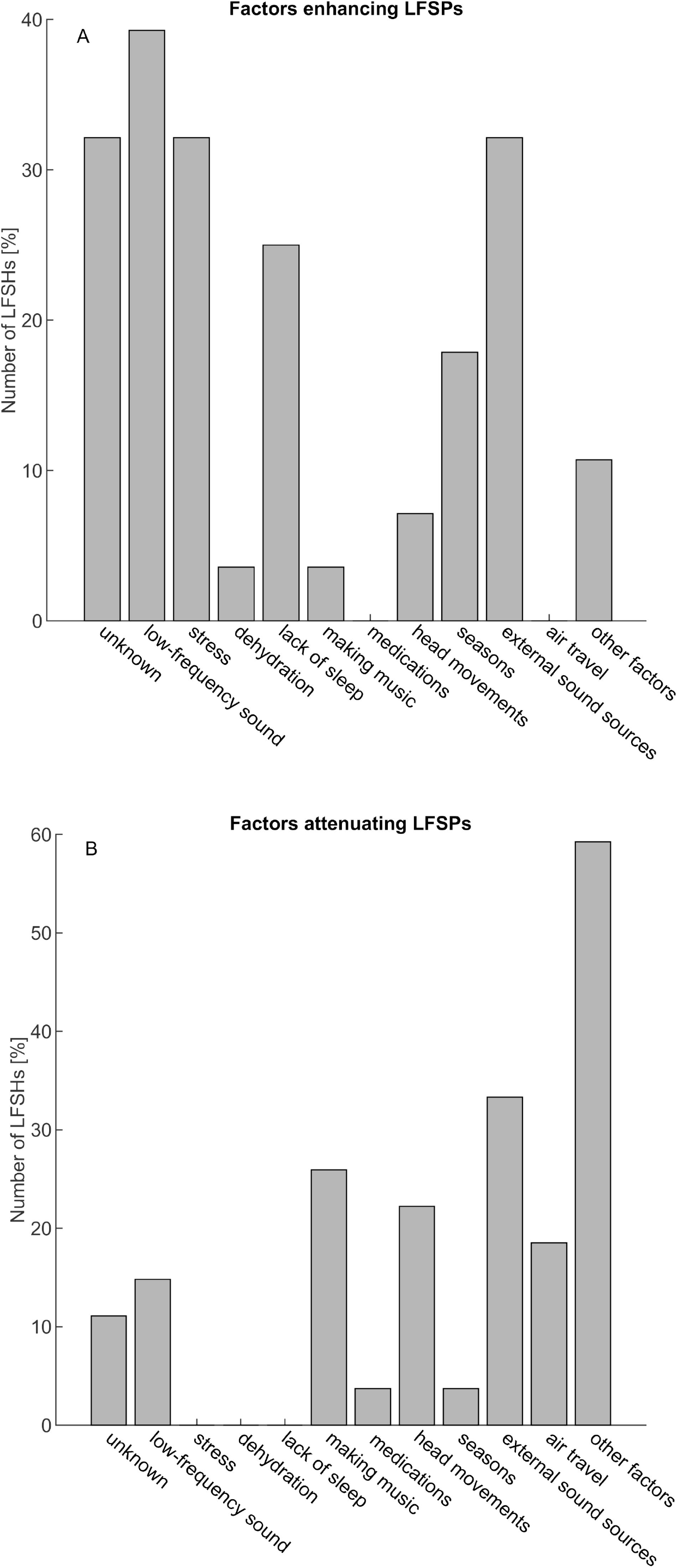
Distribution of self-reported factors enhancing (A), or attenuating (B) the LFSPs. Participants could choose several factors.

Approximately 32% of the participants reported being unaware of any factors that enhance LFSPs. Others, with a similar percentage, respectively, mentioned that low-frequency sounds, stress, or external sound sources could enhance LFSPs. Additionally, participants were given the opportunity to define their own influencing factors, which were then summarized as ‘other factors’. Here, weather conditions as enhancing factors and various activities such as travelling, outdoor activities, alcohol consumption, leisure activities or sports, and loud music as attenuating factors were mentioned. The most frequently selected options (ca. 60% of the participants) for attenuating factors were individual ‘other factors’, but around 30% of the participants also named pre-defined mitigating factors such as ‘making music’, ‘head movements’, ‘external sound sources’ and ‘air travel’.

### Low-frequency hearing thresholds

Considering the anticipated variability in low-frequency hearing thresholds resulting from the methodological approaches employed, data from a control group (CG1) was incorporated to establish a normal reference range (RR, see Fig 3). The data from the control group was collected using the same configuration and methodologies as with the LFSH group. The thresholds obtained with six default test tone frequencies (see methods) were fitted with a third-order polynomial and taken as the normal hearing threshold (NHT). The LFHTs of the LFSHs were then compared to this NHT and RR (Fig 3A). 48% of all LFHTs were lower than the interpolated NHT, and consequently 52% above the NHT, indicating no significant clustering above or below the NHT. 5% of LFHTs were lower than the RR, and 6% of LFHTs were higher than the RR, or, in other words, 89% of the LFHTs were within the RR. Some threshold values merit particular attention. Participants BT02 and BT025 presented thresholds approximately 20 dB and 30 dB below the 2.5th percentile, respectively (at 28 Hz and 62 Hz, respectively; see Fig 3A). This finding suggests that these individuals are unusually sensitive. In contrast, participant BT04 presented an outlier threshold (at 79 Hz) ca. 25 dB above the RR, being an example of unusual insensitivity at that frequency.

**Fig 3:**
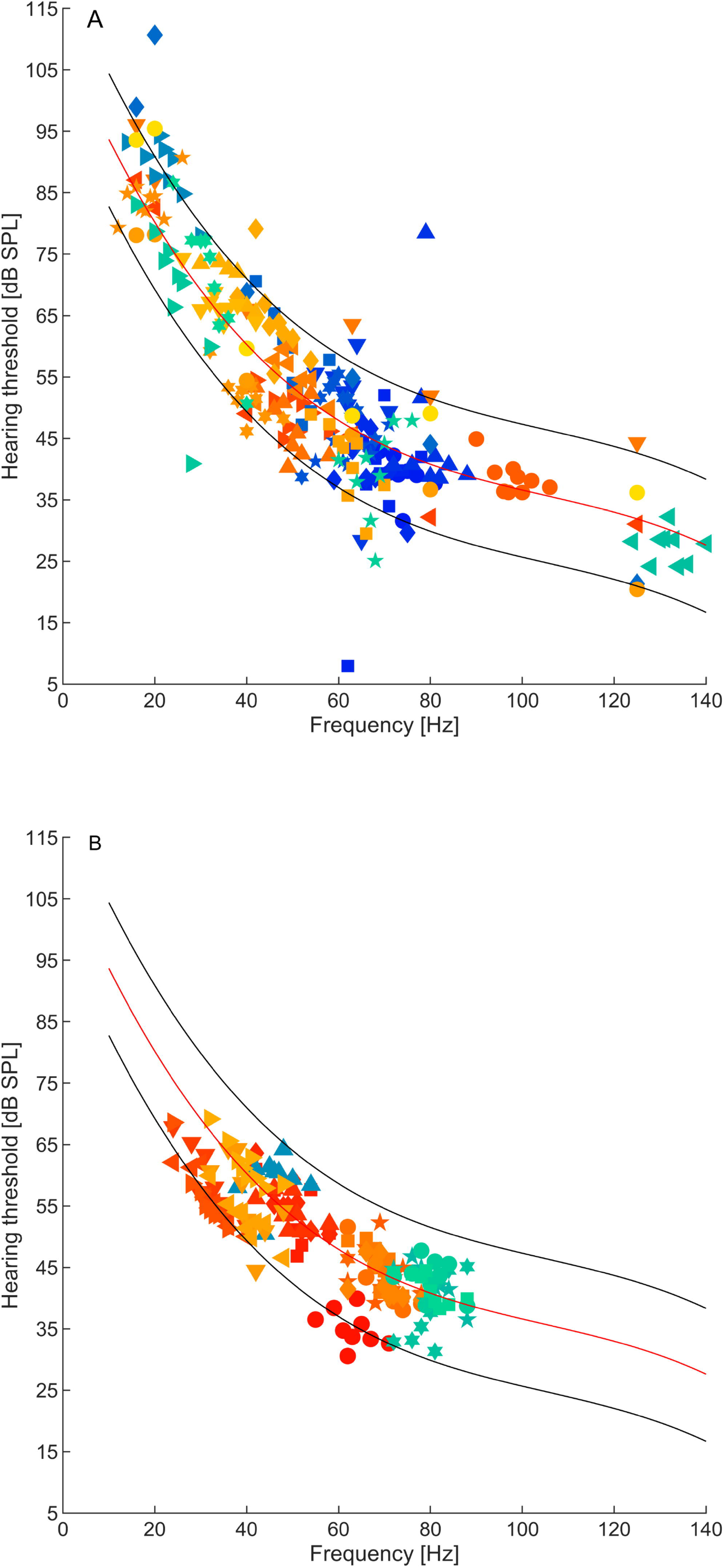
Low-frequency hearing thresholds of 28 LFSHs (A) and 7 participants from CG2 for comparison (B). Note that participants from CG2 contributed 20 hearing thresholds in total, as they were measured several times. Different marker styles or colours represent thresholds from individual participants. Thresholds from the LFSHs (A) were measured at the self-reported SPF and at additional 8 frequencies around it (for 23 LFSHs, where one LFSH completed only measurements with 6 frequencies), or, if the SPF was not available, at the 6 standard test tone frequencies (5 LFSHs, see methods). For comparison, thresholds from seven CG2 participants (B) were measured at randomly chosen, individual sets of frequencies from the LFSH group. The red line represents a polynomial fit to the low-frequency hearing thresholds obtained from a larger set of subjects belonging to another control group (CG1, N=31) that also did not report LFSPs. These reference thresholds were obtained with a standard set of frequencies. The lower and upper black lines represent the boundaries of the RR, giving the 2.5^th^ and 97.5^th^ percentile of the difference between the individual threshold values of CG1 and the polynomial fit, respectively.

Seven participants from another control group (CG2) were also tested for comparison with randomly chosen sets of test tone frequencies from the LFSH group (since a LFSP does not apply to them; see Fig 3B). Most of these thresholds (89%) fall within the established reference range.

On closer examination of the high-resolution LFHTs, threshold microstructures could be observed within both LFSHs and CG2, occurring between closely spaced test tone frequencies near the SPF (see Fig 4 for an example from a participant belonging to the LFSH group). The microstructure depth, quantified as the difference between threshold minima and maxima, showed a tendency, albeit non-significant, to be larger in LFSHs than in CG2 (medians: 16.3 dB (LFSH) and 12.6 dB (CG1), respectively, p=0.06, z-value: 1.8871, rank sum: 584).

**Fig 4:**
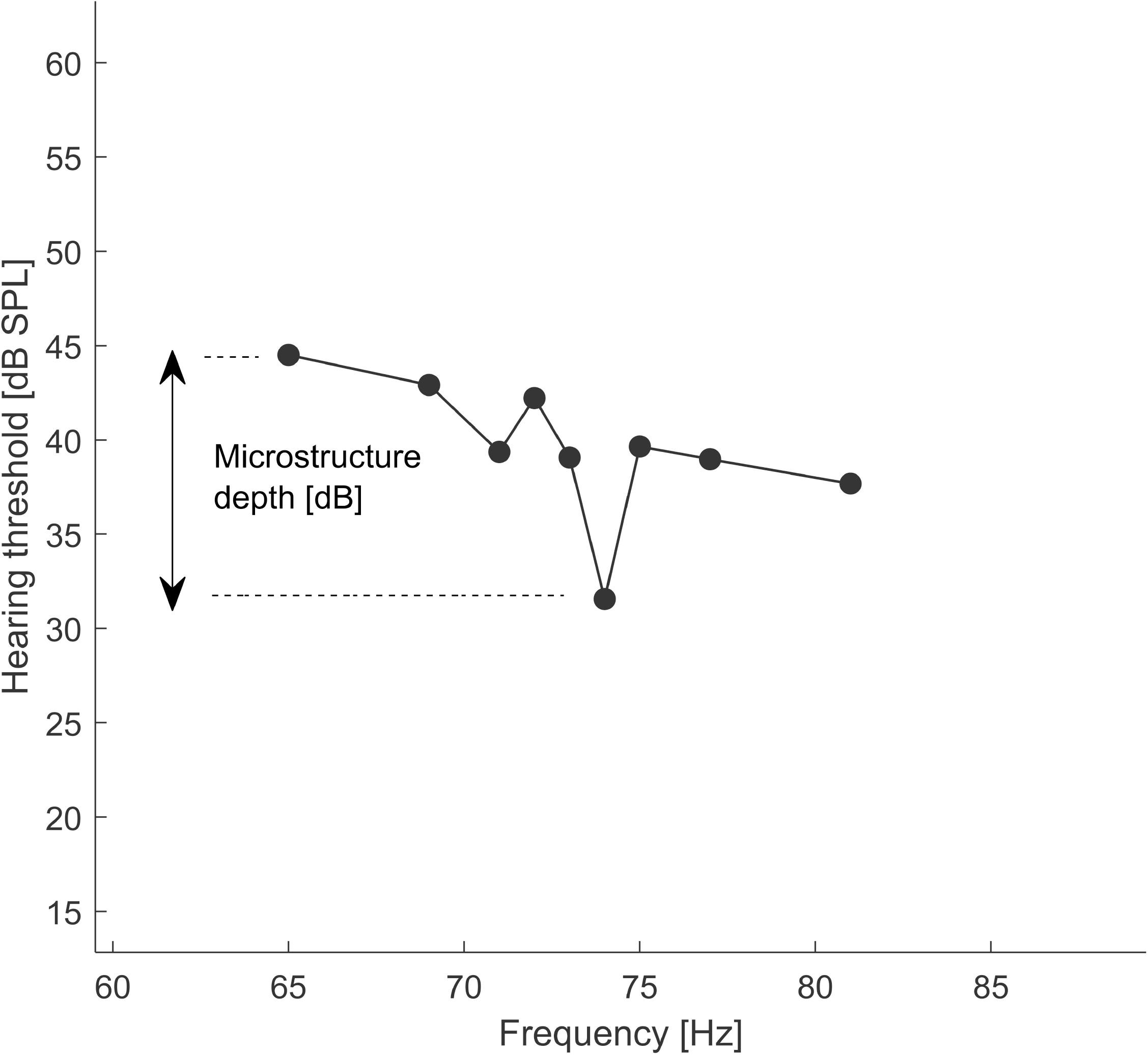
Representative example of hearing threshold microstructure from one participant in the LFSH group (BT 01), who matched the SPF to 73 Hz. The difference between the maximum and the minimum is the microstructure depth (in dB).

The distributions of individual threshold maxima and minima around the tested frequencies, expressed relative to the SPFs showed no significant difference (two-sample Kolmogorov-Smirnov test; D=0.151, p=0.68) between the participants from the LFSH group (see Fig 5A), and those in CG2 (Fig 5B), indicating no specific difference in clustering of threshold minima and maxima.

**Figure 5.**
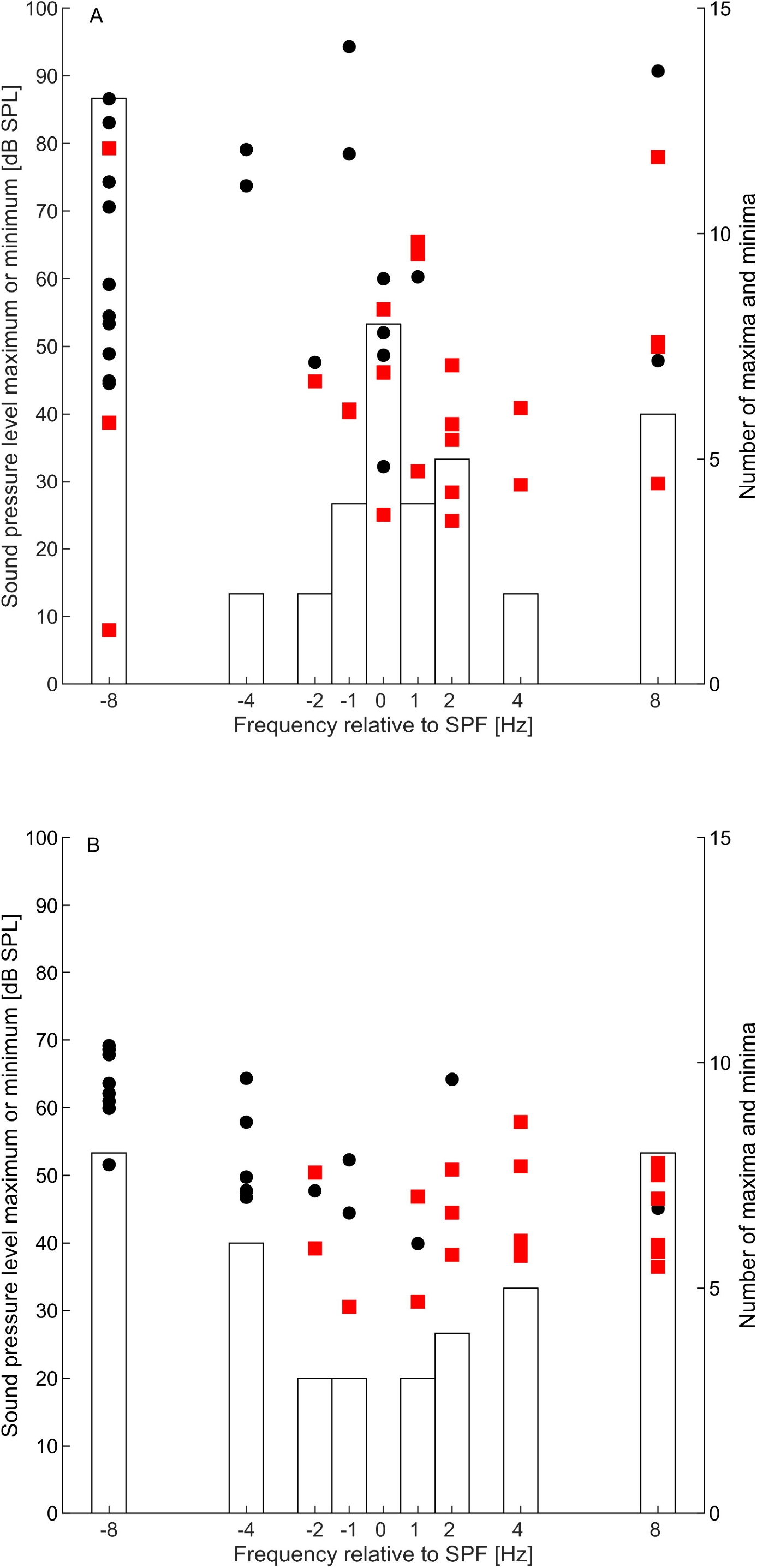
Microstructure of the hearing thresholds, expressed as a threshold maximum (black circles) and threshold minima (red squares) as a function of test tone frequency. Test tone frequencies are expressed relative to the individual SPF. The histograms represent the combined number of maxima and minima in relation to the test tone frequency. Results are shown for the LFSP group (A) and control group (CG2, B). For comparative purposes, randomly selected series of test tone frequencies from the LFSH group were selected and tested with participants belonging to CG2 (B).

### Spontaneous otoacoustic emissions

SOAEs were measured with two different OAE probes with the intention to maximize the number of SOAE candidates and to reduce the noise floor, especially in the low-frequency region, thereby facilitating the detection of low-frequency SOAEs. The employment of a dual-microphone probe and the subsequent calculation of the cross-spectral density based on the two microphone signals clearly improved the noise floor (and thus the SNR) in comparison with conventional OAE probes (Fig 6).

**Fig 6:**
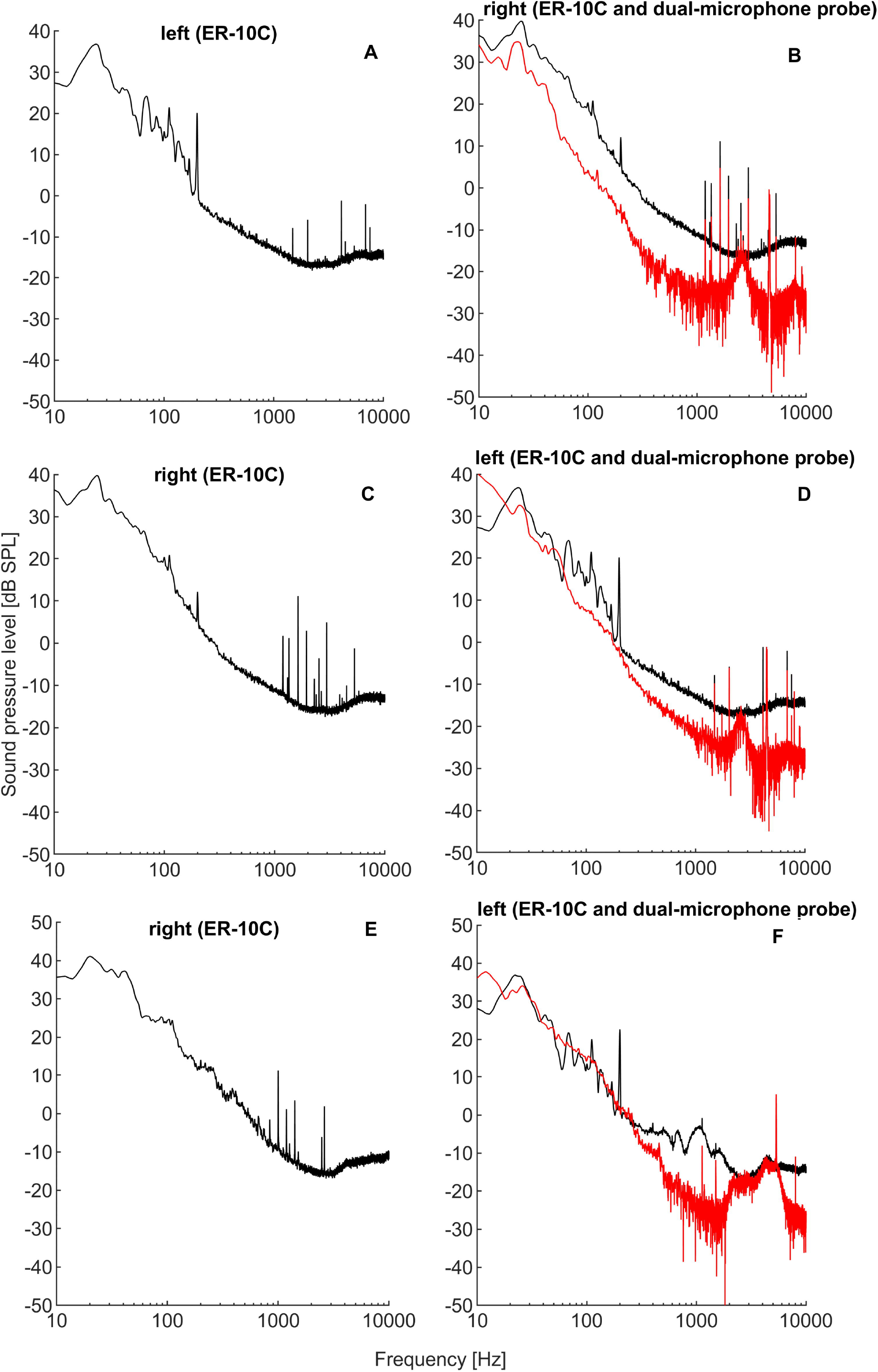
Representative examples of SOAE spectra measured in both ears of three participants (shown in the three rows, respectively) from the control groups. Here, SOAE spectra shown in the right panel (B, D, F) were successively measured with two different OAE measurement systems (black lines: ER-10C, red lines: dual-microphone probe system), while the measurements shown in the left panel (A, C, E) were only measured with the ER-10C system (black lines). The ER-10C measurements presented were measured simultaneously in both ears, as indicated above plots Employing a dual-microphone probe and a spectral cross-correlation procedure resulted in a reduced noise floor and an improvement of the SNR. Note that spectral lines with identical frequencies (within a margin of 10 Hz) in the left and right ears of the participants were considered artefacts and were omitted from the final analysis.

SOAE frequencies measured in the CG1 and LFSH groups ranged from 861 Hz to 4637 Hz (Fig 7), indicating that SOAEs with frequencies lower than about 900 Hz are generally seldom to be found or at least rarely exceed sufficient SNR using available systems. As both ears were measured in 12 tested participants of CG1, and all LFSH participants who perceived the LFSP binaurally (in some cases several times to obtain adequate measurement results with the lowest possible background noise level), all measurable SOAEs were pooled from the respective test subjects. It was found that the SOAEs of both the LFSHs and CG1 were slightly undulating in frequency, which might be caused by factors external to the ear, such as body temperature [43] or changes in body position [44], which can be regarded as physiological. SOAEs deviating by a few Hz in repeated measurements were counted as one SOAE. In the participants from CG1, SOAEs were detected in 8 out of 12 participants (67%), while in the LFSH group, this was the case for 8 out of 28 participants (29%). A total of 20 SOAEs were measured in CG1 and 17 in the LFSH group.

**Fig 7:**
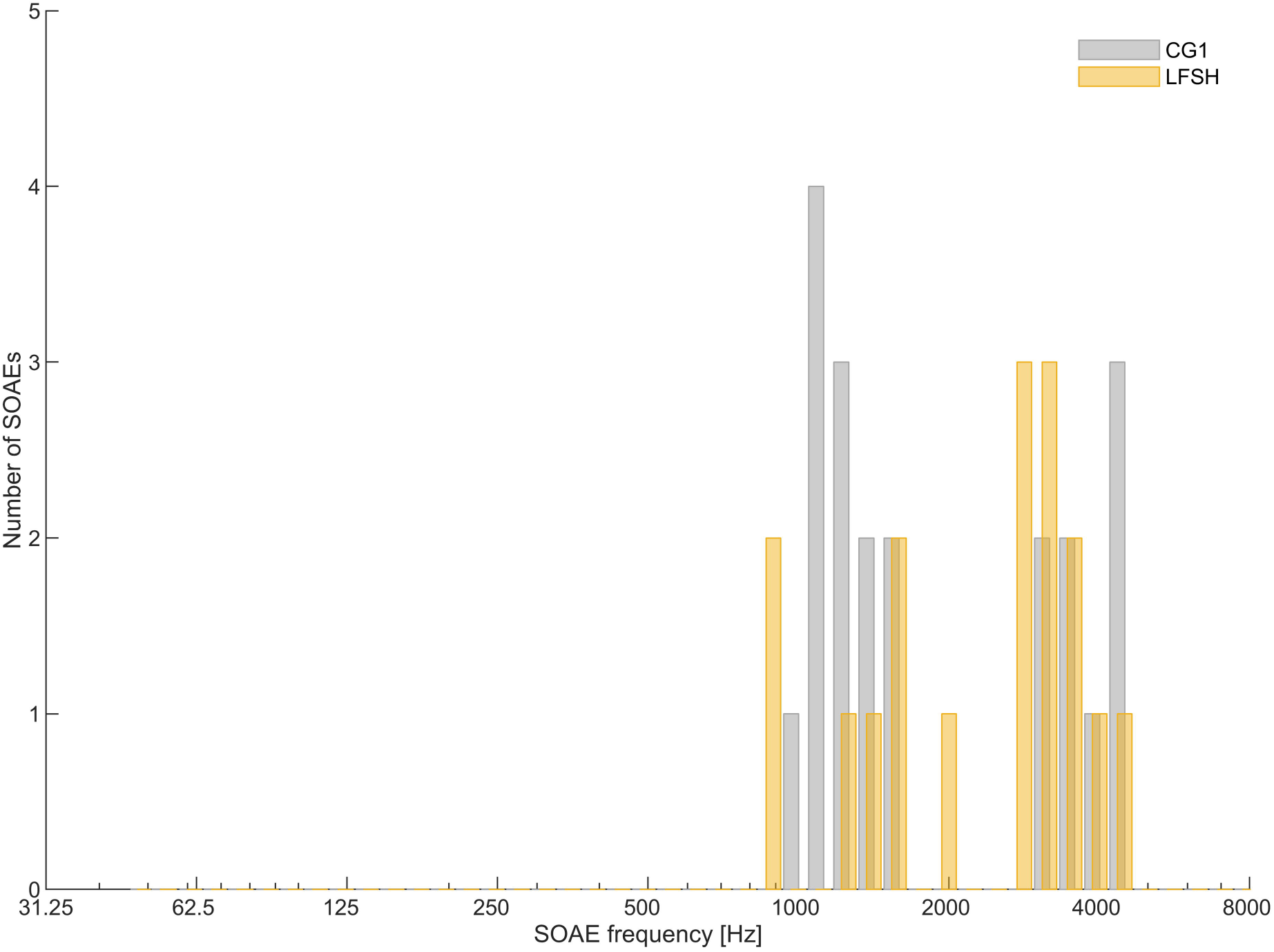
Number and frequency of valid SOAEs from the LFSH group (yellow columns) and CG1 (grey columns), bin size= 200 cents). Note the absence of SOAEs with frequencies below ca. 900 Hz. SOAE candidates above 5 kHz were not included in the analysis.

## Discussion

The present study was conducted with the objective of testing two hypotheses concerning the perception of low-frequency sounds that appear to be perceivable only by a limited number of individuals and are commonly referred to as the “Hum phenomenon”. At first glance, the observation that some complainants hear a physical sound source while others in their vicinity do not, suggests an unusually low LFHT in the relevant range. The hearing thresholds in the normal frequency range of the participants in this study agreed with their age, showing the usual signature of age-related hearing loss [45]. But normal audiometry with its frequency increments in octaves (or fractions thereof) does neither consider very low test-tone frequencies nor the microstructure of hearing thresholds, a term summarising non-monotonic, quasi-periodic hearing threshold changes over a narrow range of test tone frequencies [46]. These highly individual peaks and troughs in the hearing thresholds might at least represent good candidates for unusually low hearing thresholds, and microstructures of LFHTs have been demonstrated before [16]. In other words, there is a good chance that regions with unusually low thresholds in the low-frequency range of hearing are being overlooked by conventional audiometry.

Measuring threshold microstructure is time-consuming and strenuous for the participants and therefore were test tone frequencies in this study limited to a very narrow frequency range, with very small frequency increments and decrements around the self-tested SPF. To limit the number of frequency steps required, frequency steps were increased with increasing distance from the SPF. The observed increased fine structure around the SPF could hence be the result of tighter “sampling”, corroborated by the fact that the control groups show a comparable microstructure. Most of the measured thresholds fall within the central 95% of the NHTs (i.e. the range from the 2.5^th^ to the 97.5^th^ percentile) that we measured in a young, normal-hearing, non-LFSP-hearing cohort. Assuming that the LFSHs are indeed characterised by unusually low thresholds in the low-frequency range of hearing, it would be expected that a significant proportion of the measured thresholds are below the 2.5^th^ percentile, the lower limit of the reference range, or at least in the lower half of the threshold distribution. This was, with the notable exception of two thresholds from two participants, not the case. It is nonetheless not unlikely that in those cases of LFSPs where a physical sound source can be assumed to be underlying, it is simply a problem of normality, as threshold differences of up to about 20 dB can be expected between the most sensitive and least sensitive in a *normal* population [13]. In future studies, those in the familial units of LFSHs not hearing a LFSP should be included in the testing. The thresholds measured here concur with those reported in [17], were also individuals experiencing LFSPs were tested, and where in general no abnormally sensitive thresholds were found. As the authors of that study did not attempt to measure threshold microstructure, the possibility was raised that local threshold minima might not have been detected. Here, we showed that this is indeed the case, but threshold microstructure does not seem to be sufficient to explain LFSPs, as the control group in our study also showed comparable threshold microstructure, without reporting LFSPs. Still, present results suggest that within the small percentage of LFSP complainants in relation the overall population, there is an even smaller percentage of individuals that do present abnormally high sensitivity. The latter has been observed before (discussed in [10]), and might stem from anatomical variations in the inner ear. For example, intracochlear pressure produced by LFS been found to largely influenced by the helicotrema, which is an opening in the cochlear apex of the inner ear that shunts pressure differences across the cochlear fluid-filled scalae and thus decreases the inner ear’s sensitivity for LFS. It has been found that dips and peaks in the intracochlear pressure are an expected effect of the helicotrema impedance, and that these are reflected in both hearing threshold and loudness perception for sounds below 100 Hz [16, 47]. Thus, anatomical variations in this structure might render a very small percentage of individuals more sensitive than normal.

Participants in this study included both complainants who reported that a LFS source could be identified, and those who could not. About 2/3 of LFSHs in this study reported that it was only them who heard the LFS. Thus, the LFSHs in this study may have involved both cases of unusually low LFHT, and cases with normal LFHTs where possibly an aversion to these sounds was developed. The LFHTs measured in this study suggest that both case types were present, albeit the majority of LFSHs presented normal LFHTs and only a few could be classified as especially sensitive.

The second hypothesis explored in this study suggests that the perception of LFSPs is due to hearing one’s own spontaneous otoacoustic emissions in the low-frequency range. However, SOAEs are typically not audible to their owners, despite sound levels which may exceed the hearing thresholds for that frequency [21]. Additionally, low-frequency SOAEs can be difficult to measure, and in the present study these were not detected at frequencies below about 900 Hz in either the LFSH group or the control group. This finding is consistent with previous research on SOAE frequencies in the normal population. It is, however, unclear if unsuccessful attempts have been previously made to measure SOAEs in the low-frequency range, or whether the lack of reported SOAE frequencies in this range has been due to arbitrarily restricting the search for candidate SOAEs to frequencies higher than ca. 500 Hz [21]. Since the absence of SOAEs with frequencies below about 900 Hz could simply indicate that the SNR was not sufficiently large at low frequencies, we attempted to improve this by using dedicated equipment and signal processing. Despite these efforts, no SOAEs with frequencies below approximately 900 Hz could be found in the present study. It seems that, based on our results, two rare events need to coincide for SOAEs to serve as an explanation for a LFSP: the occurrence of supra-threshold, low-frequency SOAE, and their audibility. This renders hearing low-frequency SOAEs an unlikely explanation for LFSPs.

Potential low frequency oscillations from the peripheral hearing system are not only limited to SOAEs: it has been known for several years that the tympanic membranes move during eye movements (eye movement related eardrum oscillations, EMREOs), and those, presumably caused by middle ear muscle activity, can be registered with microphones in the ear canal [56–59], similar to SOAEs. The recorded pressure changes have a period length corresponding to a frequency of ca. 30 Hz, with a total duration of ca. 100 ms, typically occurring 3 times/ second during saccadic eye movements [56], and therefore resembling most likely a rhythmic sensation if audible. With reported levels of ca. 60 peak-equivalent dB SPL, typical EMREOs are most likely inaudible at the frequencies involved. Assuming some LFSHs can hear EMREOs, the postulated rhythmic percept is not in good agreement with reports on the nature of LFSPs, making EMREOs an unlikely explanation for LFSPs.

It has also been shown that attention to visual stimuli induces very low-frequency amplitude modulations of cubic distortion product otoacoustic emissions (the carrier) [60], but it is unclear if the observed modulation depth is large enough to be perceived (assuming that external sound stimuli undergo a similar amplitude modulation during visual attention). If it is perceivable, the amplitude modulation is unlikely to lead to the tonal percepts often reported by LFSHs [17].

Finally, although not directly tested in this study, low-frequency tinnitus is a plausible explanation for LFSPs in those with normal hearing thresholds, and where no ESS can be identified. However, complainants might consider this unlikely, maybe because they externalise the LFSP they hear, which further complicates the consideration of the possibility that what they hear is a tinnitus-like phenomenon, when no external sound sources can be verified. Externalisation is the natural percept one experiences with real, external sound sources, whereas internalisation is the “unnatural” percept related for example to listening to music via conventional headphones, and perceiving the music “inside the head”. Th internalisation of a sound can be thought of as the consequence of the absence and/ or violation of binaural and other cues expected from a real sound source, compared to the cues available from an internal tinnitus source (or conventional headphones, for that matter). This includes the absence of spectral cues [48], reverberation [49], and changing binaural cues with head movements [48, 50, 51]. Externalisation, on the other hand, occurs if those cues are available and realistic [51, 52]. While some cues are available to humans when it comes to the localisation of low-frequency sound sources, minimum audible angles are much larger compared to sound sources in the normal frequency range, even more so in a room small enough to have dominant room modes [53]. Since available natural cues for localising low-frequency sound sources in a room are quite rudimental, even an internal generator of a low-frequency tinnitus might satisfy the conditions needed to externalise the percept. Reports about subjective tinnitus (albeit non low-frequency) with perceived spatial localisations outside of the head exist [54]. For those suffering from a low-frequency tinnitus percept, retraining strategies involving alternating listening to artificial sound sources presented in the sound field and within the ear canal might facilitate the process of internalisation, normally occurring in those experiencing subjective tinnitus [54].

We conclude that hearing threshold hypersensitivity in the low-frequency range can only serve as an LFSP explanation for very few LFSHs, and while not directly tested in this study, low-frequency tinnitus might serve as a good explanation for many, but not all, cases. No low-frequency SOAEs were detected in neither controls groups nor LFSHs, ruling out audible SOAEs as the source. Although the substantially larger background noise in the low-frequency range makes detection of SOAEs more difficult, audible SOAEs in the low-frequency can be expected to have large amplitudes, because of the hearing thresholds involved in this frequency range. It appears that LFSPs cannot be explained by a single mechanism but may be the result of sources that vary from individual to individual. As outlined before, this study was limited to auditory explanations regarding the LFSPs, but this does not rule out that other explanations, rooted in other scientific disciplines, are not plausible.

## Supporting information

Hearing thresholds CG1

Hearing thresholds CG2

Hearing thresholds LFSH

Questionnaire results

SOAES CG1

SOAEs LFSH

Questionnaire german

Questionaire english translation

## Acknowledgements

The authors would like to thank all participants for their contributions to this study.

## Supporting information

**S1 data hearing thresholds control group 1 (G1)**

**S2 data hearing threshold control group 2 (CG2)**

**S3 data low frequency hearing thresholds (LFSH) S4 questionnaire results**

**S5 data spontaneous otoacoustic emission (SOAES) from control group 1 (CG1)**

**S6 data spontaneous otoacoustic emission (SOAES) from control group 1 (LFSH)**

**S7 questionnaire in German**

**S8 questionnaire English translation**

## References

1. Frosch FG. Manifestations of a low-frequency sound of unknown origin perceived worldwide, also known as “the Hum” or the “Taos Hum”. Int Tinnitus J. 2016;20(1):59–63. Epub 20160722. doi: 10.5935/0946-5448.20160011. PubMed PMID: 27488996.

2. Leventhall HG. Low frequency noise and annoyance. Noise Health. 2004;6(23):59–72. PubMed PMID: 15273024.

3. Årdal OK. Hege høyrer noko nesten ingen andre kan høyre. nrk. 2022. Available from: https://www.nrk.no/kultur/xl/the-hum_-hege-hoyrer-noko-nesten-ingen-andre-hoyrer-1.16121231.

4. Joachim J. Dem mysteriösen „Brummton-Phänomen” auf der Spur. Frankfurter Rundschau. 2021. Available from: https://www.fr.de/rhein-main/darmstadt/darmstadt-dem-brummton-phaenomen-auf-der-spur-91055609.html.

5. Sheik I. “Windsor Hum”: The mystery noise that plagued thousands of Canadians for years — until the pandemic ended it. DH news. 2022. Available from: https://dailyhive.com/vancouver/windsor-hum-ontario.

6. MacPherson D. The World Hum Map and Database Project. Available from: https://www.thehum.info/.

7. Hessisches Landesamt für Naturschutz UuG. Messbericht über Geräuschmessungen zur Ermittlung von tieffrequenten Geräuschen in Darmstadt (Martinsviertel). 2011. Available from: https://www.darmstadt.de/fileadmin/PDF-Rubriken/Leben_in_Darmstadt/umwelt/Laerm/Messbericht__2021.pdf

8. Leventhall G. Still Humming. Noise Vibration Worldwide. 2005;36(2):21–6. doi: 10.1260/0957456053499103.

9. Leventhall G. Low Frequency Noise. What we know, what we do not know, and what we would like to know. Journal of Low Frequency Noise, Vibration and Active Control. 2009;28(2):79–104. doi: 10.1260/0263-0923.28.2.79.

10. Moller H, Pedersen CS. Hearing at low and infrasonic frequencies. Noise Health. 2004;6(23):37–57. PubMed PMID: 15273023.

11. Berglund B, Hassmen P, Job RF. Sources and effects of low-frequency noise. J Acoust Soc Am. 1996;99(5):2985–3002.

12. Leventhall G, Pelmear P, Benton S. A Review of Published Research on Low Frequency Noise and its Effects 2003 [updated 01/01]. Available from: https://webarchive.nationalarchives.gov.uk/ukgwa/20081106144101/ http://www.defra.gov.uk/environment/noise/research/lowfrequency/pdf/lowfreqnoise.pdf.

13. Kurakata K, Mizunami T, Matsushita K. Percentiles of normal hearing-threshold distribution under free-field listening conditions in numerical form. Acoust Sci Technol. 2005;26(5):447–9. doi: 10.1250/ast.26.447.

14. Long GR. The microstructure of quiet and masked thresholds. Hear Res. 1984;15(1):73–87. doi: 10.1016/0378-5955(84)90227-2.

15. Dewey JB. Sources of Microstructure in Mammalian Cochlear Responses. JARO. 2025. doi: 10.1007/s10162-025-00974-5.

16. Frost GP. An Investigation into the Microstructure of the Low Frequency Auditory Threshold and of the Loudness Function in the near Threshold Region. Journal of Low Frequency Noise, Vibration and Active Control. 1987;6(1):34–9. doi: 10.1177/026309238700600104.

17. Pedersen CS, Møller H, Waye KP. A Detailed Study of Low-Frequency Noise Complaints. Journal of Low Frequency Noise, Vibration and Active Control. 2008;27(1):1–33. doi: 10.1260/026309208784425505.

18. Luecke VN, Buchwieser L, Eulenburg Pz, Marquardt T, Drexl M. Ocular and cervical vestibular evoked myogenic potentials elicited by air-conducted, low-frequency sound. Journal of Vestibular Research-Equilibrium & Orientation. 2020;30(4):235–47. doi: 10.3233/ves-200712. PubMed PMID: WOS:000582717000002.

19. Rajala V, Hakala J, Alakoivu R, Koskela V, Hongisto V. Hearing threshold, loudness, and annoyance of infrasonic versus non-infrasonic frequencies. Applied Acoustics. 2022;198:108981. doi: 10.1016/j.apacoust.2022.108981.

20. Yamada S, Ikuji M, Fujikata S, Watanabe T, Kosaka T. Body Sensation of Low Frequency Noise of Ordinary Persons and Profoundly Deaf Persons. Journal of Low Frequency Noise, Vibration and Active Control. 1983;2(3):32–6. doi: 10.1177/026309238300200302.

21. Probst R, Lonsbury-Martin BL, Martin GK. A review of otoacoustic emissions. J Acoust Soc Am. 1991;89(5):2027–67. doi: 10.1121/1.400897. PubMed PMID: 1860995.

22. Shera CA. Whistling While it Works: Spontaneous Otoacoustic Emissions and the Cochlear Amplifier. Journal of the Association for Research in Otolaryngology: JARO. 2022;23(1):17–25. Epub 20220103. doi: 10.1007/s10162-021-00829-9. PubMed PMID: 34981262; PubMed Central PMCID: PMCPMC8782959.

23. Zurek PM. Spontaneous narrowband acoustic signals emitted by human ears. J Acoust Soc Am. 1981;69(2):514–23. Epub 1981/02/01. PubMed PMID: 7462474.

24. Strickland EA, Burns EM, Tubis A. Incidence of spontaneous otoacoustic emissions in children and infants. J Acoust Soc Am. 1985;78(3):931–5. doi: 10.1121/1.392924. PubMed PMID: 4031263.

25. Long G. Perceptual consequences of the interactions between spontaneous otoacoustic emissions and external tones. I. Monaural diplacusis and aftertones. Hear Res. 1998;119(1-2):49–60. doi: Doi 10.1016/S0378-5955(98)00032-X. PubMed PMID: WOS:000073795600005.

26. Burns EM. Long-term stability of spontaneous otoacoustic emissions. J Acoust Soc Am. 2009;125(5):3166–76. Epub 2009/05/12. doi: 10.1121/1.3097768. PubMed PMID: 19425659; PubMed Central PMCID: PMCPmc2806441.

27. Fritze W. Registration of spontaneous cochlear emissions by means of Fourier transformation. Arch Otorhinolaryngol. 1983;238(2):189–96. doi: 10.1007/BF00454312. PubMed PMID: 6626031.

28. Mcfadden D, Wightman FL. Audition - Some Relations between Normal and Pathological Hearing. Annu Rev Psychol. 1983;34:95–128. doi: DOI 10.1146/annurev.ps.34.020183.000523. PubMed PMID: WOS:A1983QA76200005.

29. Penner MJ. Linking spontaneous otoacoustic emissions and tinnitus. Br J Audiol. 1992;26(2):115–23. doi: 10.3109/03005369209077879.

30. Henry JA, Dennis KC, Schechter MA. General review of tinnitus: Prevalence, mechanisms, effects, and management. J Speech Lang Hear R. 2005;48(5):1204–35. doi: 10.1044/1092-4388(2005/084). PubMed PMID: WOS:000234963100018.

31. Lockwood AH, Salvi RJ, Burkard RF. Tinnitus. N Engl J Med. 2002;347(12):904–10. doi: 10.1056/NEJMra013395. PubMed PMID: 12239260.

32. Ciocon JO, Amede F, Lechtenberg C, Astor F. Tinnitus - a Stepwise Work-up to Quiet the Noise Within. Geriatrics. 1995;50(2):18–25. PubMed PMID: WOS:A1995QF59800005.

33. Baguley D, McFerran D, Hall D. Tinnitus. The Lancet. 2013;382(9904):1600–7. doi: 10.1016/S0140-6736(13)60142-7.

34. Haider HF, Bojic T, Ribeiro SF, Paço J, Hall DA, Szczepek AJ. Pathophysiology of Subjective Tinnitus: Triggers and Maintenance. Front Neurosci. 2018;12. doi: ARTN 866 10.3389/fnins.2018.00866. PubMed PMID: WOS:000451467900001.

35. Shore S, Zhou JX, Koehler S. Neural mechanisms underlying somatic tinnitus. Tinnitus: Pathophysiology and Treatment. 2007;166:107–+. doi: 10.1016/S0079-6123(07)66010-5. PubMed PMID: WOS:000280613600012.

36. Meikle M, Taylor-Walsh E. Characteristics of tinnitus and related observations in over 1800 tinnitus clinic patients. J Laryngol Otol Suppl. 1984;9:17–21. doi: 10.1017/s1755146300090053. PubMed PMID: 6596358.

37. Walford RE. A Classification of Environmental “Hums” and Low Frequency Tinnitus. Journal of Low Frequency Noise, Vibration and Active Control. 1983;2.

38. Kuehler R, Fedtke T, Hensel J. Infrasonic and low-frequency insert earphone hearing threshold. J Acoust Soc Am. 2015;137(4):EL347–53. Epub 2015/04/30. doi: 10.1121/1.4916795. PubMed PMID: 25920888.

39. Berg D. SoundMexPro - The free professional Tool for multi-channel I/O and real-time Processing of Audio Signals in MATLAB®, GNU Octave and Python. Zenodo; 2024.

40. Braun M. Accurate binaural mirroring of spontaneous otoacoustic emissions suggests influence of time-locking in medial efferents. Hear Res. 1998;118(1):129–38. doi: 10.1016/S0378-5955(98)00028-8.

41. Kugler K, Wiegrebe L, Grothe B, Kössl M, Gürkov R, Krause E, et al. Low-frequency sound affects active micromechanics in the human inner ear. Royal Society Open Science. 2014;1(2).

42. Maat B, De Kleine E, Van Dijk P. Spontaneous otoacoustic emission measurement with dual microphone technique ARO 39th MidWinter Meeting; Manchester Grand Hyatt, San Diego, California, USA 2016.

43. Wit HP. Diurnal cycle for spontaneous oto-acoustic emission frequency. Hear Res. 1985;18(2):197–9. doi: 10.1016/0378-5955(85)90012-7. PubMed PMID: 4044421.

44. de Kleine E, Wit HP, van Dijk P, Avan P. The behavior of spontaneous otoacoustic emissions during and after postural changes. J Acoust Soc Am. 2000;107(6):3308–16. Epub 2000/06/30. PubMed PMID: 10875376.

45. Agrawal Y, Platz EA, Niparko JK. Prevalence of Hearing Loss and Differences by Demographic Characteristics Among US Adults: Data From the National Health and Nutrition Examination Survey, 1999-2004. Arch Intern Med. 2008;168(14):1522–30. doi: 10.1001/archinte.168.14.1522.

46. Mauermann M, Long G, Kollmeier B. Fine structure of hearing threshold and loudness perception. The Journal of the Acoustical Society of America. 2004;116:1066–80. doi: 10.1121/1.1760106.

47. Jurado C, Marquardt T. The effect of the helicotrema on low-frequency loudness perception. The Journal of the Acoustical Society of America. 2016;140:3799–809. doi: 10.1121/1.4967295.

48. Blauert J. Spatial Hearing: The Psychophysics of Human Sound Localization: The MIT Press; 1996.

49. Plenge G. On the differences between localization and lateralization. J Acoust Soc Am. 1974;56(3):944–51. doi: 10.1121/1.1903353. PubMed PMID: 4420742.

50. Wallach H. The role of head movements and vestibular and visual cues in sound localization. J Exp Psychol. 1940;27:339–68.

51. Brimijoin WO, Boyd AW, Akeroyd MA. The Contribution of Head Movement to the Externalization and Internalization of Sounds. PLoS One. 2013;8(12):e83068. doi: 10.1371/journal.pone.0083068.

52. Hartmann WM, Wittenberg A. On the externalization of sound images. The Journal of the Acoustical Society of America. 1996;99(6):3678–88. doi: 10.1121/1.414965.

53. Hill A, Lewis SP, Hawksford MOJ, editors. Towards a generalized theory of low-frequency sound source localization. Reproduced Sound 2012; 2012.

54. Searchfield GD, J. Sp, and Barde A. A scoping review of the spatial perception of tinnitus and a guideline for the minimum reporting of tinnitus location. J R Soc N Z. 2025;55(3):501–19. doi: 10.1080/03036758.2024.2344781.

55. Potgieter I, MacDonald C, Partridge L, Cima R, Sheldrake J, Hoare DJ. Misophonia: A scoping review of research. J Clin Psychol. 2019;75(7):1203–18. Epub 20190311. doi: 10.1002/jclp.22771. PubMed PMID: 30859581.

56. Gruters KG, Murphy DLK, Jenson CD, Smith DW, Shera CA, Groh JM. The eardrums move when the eyes move: A multisensory effect on the mechanics of hearing. Proc Natl Acad Sci U S A. 2018;115(6):E1309–e18. Epub 20180123. doi: 10.1073/pnas.1717948115. PubMed PMID: 29363603; PubMed Central PMCID: PMCPMC5819440.

57. Lovich SN, King CD, Murphy DLK, Abbasi H, Bruns P, Shera CA, et al. Conserved features of eye movement related eardrum oscillations (EMREOs) across humans and monkeys. Philosophical Transactions of the Royal Society B: Biological Sciences. 2023;378(1886):20220340. doi:10.1098/rstb.2022.0340.

58. Abbasi H, King CD, Lovich S, Röder B, Groh JM, Bruns P. Eye movement-related eardrum oscillations do not require current visual input. Hear Res. 2025;465:109346. doi: 10.1016/j.heares.2025.109346.

59. King CD, Lovich SN, Murphy DLK, Landrum R, Kaylie D, Shera CA, et al. Individual similarities and differences in eye-movement-related eardrum oscillations (EMREOs). Hear Res. 2023;440:108899. doi: 10.1016/j.heares.2023.108899.

60. Dragicevic CD, Marcenaro B, Navarrete M, Robles L, Delano PH. Oscillatory infrasonic modulation of the cochlear amplifier by selective attention. PLoS One. 2019;14(1):e0208939. doi: 10.1371/journal.pone.0208939.

