## Supplementary material for "On the potential sources of a low-frequency sound percept only a few can perceive": Questionnaire german

### Fragebogen

1. Wie beurteilen sie die akustische Situation an Ihrem Wohnort auf einer Skala von 1-10?

| 1 | 2 | 3 | 4 | 5 | 6 | 7 | 8 | 9 | 10 |
|---|---|---|---|---|---|---|---|---|---|
| ruhig | | | | | | | | | laut |

2. Wann wurde der Brummton das erste Mal von Ihnen wahrgenommen (MM/JJJJ)?

_________

3. Haben Sie zusätzliche, andere akustische Wahrnehmungen unbekannter Herkunft (‚normaler' Tinnitus)?

☐ Ja
Welche:________________________________________
☐ Nein

4. Ist Ihre Brummtonwahrnehumg eindeutig einem Ohr zuordenbar?

☐ Nein
☐ Ja, dem linken Ohr
☐ Ja, dem rechten Ohr

5. Welche Faktoren verstärken Ihren Brummton?

☐ Keine
☐ Tieffrequente Schalle
☐ Stress
☐ Dehydration
☐ Schlafmangel
☐ Musizieren
☐ Medikamente
☐ Kopfbewegungen
☐ Jahreszeit

- ☐ Externe Schallquellen
- ☐ Flugreisen
- ☐ Sonstige:___________________________________________

6. Welche Faktoren verringern Ihren Brummton?

- ☐ Keine
- ☐ Tieffrequente Schalle
- ☐ Stress
- ☐ Dehydration
- ☐ Schlafmangel
- ☐ Musizieren
- ☐ Medikamente
- ☐ Kopfbewegungen
- ☐ Jahreszeit
- ☐ Externe Schallquellen
- ☐ Flugreisen
- ☐ Sonstige:___________________________________________

7. Tritt der Brummton nur an bestimmten Orten auf?

- ☐ Ja
  Welche:___________________________________________
- ☐ Nein

8. Ist es dort eher leise oder laut? Beurteilen Sie dies auf einer Skala von 1-10:

| 1 | 2 | 3 | 4 | 5 | 6 | 7 | 8 | 9 | 10 |
|---|---|---|---|---|---|---|---|---|---|
| ruhig | | | | | | | | | laut |

- ☐ unterschiedlich

9. Tritt der Brummton v.a. zu bestimmten Tageszeiten auf?

- ☐ Ja: morgens
- ☐ Ja: mittags
- ☐ Ja: abends

- ☐ Ja: nachts
- ☐ Nein

10. Nehmen weitere Familienmitglieder den Brummton wahr?

- ☐ Ja
- ☐ Nein

11. Wird der Brummton von anderen, nicht verwandten Personen in der gleichen Umgebung mit der gleichen Häufigkeit gehört?

- ☐ Ja
- ☐ Nein

12. Bestehen bei Ihnen bekannte Erkrankung des Hörsystems oder des Gleichgewichtssystems?

- ☐ Ja
- ☐ Nein

13. Nehmen Sie regelmäßig Aspirin oder vergleichbare Produkte zur Linderung der Brummtonwahrnehmung ein?

- ☐ Ja
- ☐ Nein

14. Haben seit der ersten Wahrnehmung des Brummtons Veränderungen hinsichtlich der Lautstärke des Brummtons bemerkt?

- ☐ Ja: lauter
- ☐ Ja: leiser
- ☐ Nein

15. Haben seit der ersten Wahrnehmung des Brummtons Veränderungen hinsichtlich der Tonhöhe des Brummtons bemerkt?

- ☐ Ja: tiefer
- ☐ Ja: höher
- ☐ Nein

16. Haben seit der ersten Wahrnehmung des Brummtons Veränderungen hinsichtlich der Häufigkeit des Brummtons bemerkt?

- ☐ Ja: häufiger
- ☐ Ja: seltener
- ☐ Nein

17. Hat eine Messung am Wohnort das Vorhandensein einer physikalisch nachweisbaren Tieftonschallquelle bestätigt?

- ☐ Ja
- ☐ Nein
- ☐ Nicht vorgenommen

18. Stellt die Wahrnehmung des Brummtons für Sie eine physiologische oder psychologische Belastung dar?

- ☐ Ja
- ☐ Nein

19. Worin vermuten Sie die Ursache für Ihre Wahrnehmung des Brummtons?
Externe Schallquelle

- ☐ Tinnitus
- ☐ Spontane otoakustische Emissionen
- ☐ Sonstiges:___________________________________________

20. Wie würden Sie die Qualität Ihrer Brummtonwahnehmung beschreiben?

- ☐ Reintonhaft
- ☐ Pulsierend
- ☐ An- und abschwellend
- ☐ Konstant

21. Wie alt sind Sie?

_________

22. Welches Geschlecht haben Sie?

- ☐ Männlich
- ☐ Weiblich
- ☐ Divers
