## Supplementary material for "On the potential sources of a low-frequency sound percept only a few can perceive": Questionaire english translation

Questionnaire

1. how would you rate the acoustic situation at your place of residence on a scale of 1-10?

1 2 3 4 5 6 7 8 9 10

quiet loud

2. when did you hear the humming sound for the first time (MM/YYYY)?

_________

3. do you have additional, other acoustic perceptions of unknown origin ('normal' tinnitus)?

☐ Yes

Which:_______________________________________

☐ No

4. can your hum perception be clearly assigned to one ear?

☐ No

☐ Yes, the left ear

☐ Yes, the right ear

5. which factors amplify your humming sound?

☐ None

☐ Low-frequency sounds

☐ Stress

☐ Dehydration

☐ Lack of sleep

☐ Making music

☐ Medication

☐ Head movements

☐ Season

☐ External sound sources

☐ Air travel

☐ Other:________________________________________

6. what factors reduce your hum?

☐ None

☐ Low-frequency noise

☐ Stress

☐ Dehydration

☐ Lack of sleep

☐ Making music

☐ Medication

☐ Head movements

☐ Season

☐ External sound sources

☐ Air travel

☐ Other:________________________________________

7. does the humming sound only occur in certain places?

☐ Yes

Which:________________________________________

☐ No

8. is it rather quiet or loud there? Rate this on a scale of 1-10:

1 2 3 4 5 6 7 8 9 10

quiet loud

☐ different

9. does the humming sound occur mainly at certain times of the day?

☐ Yes: in the morning

☐ Yes: at midday

☐ Yes: in the evening

☐ Yes: at night

☐ No

10. do other family members hear the humming sound?

☐ Yes

☐ No

11. is the humming sound heard by other, unrelated people in the same neighbourhood with the same frequency?

☐ Yes

☐ No

12. do you have any known illnesses of the hearing system or the balance system?

☐ Yes

☐ No

13. do you regularly take aspirin or similar products to alleviate the perception of buzzing?

☐ Yes

☐ No

14. have you noticed any changes in the volume of the humming sound since you first heard it?

☐ Yes: louder

☐ Yes: quieter

☐ No

15. have you noticed any changes in the pitch of the humming sound since you first heard it?

☐ Yes: lower

☐ Yes: higher

☐ No

16. have you noticed any changes in the frequency of the humming sound since you first heard it?

□ Yes: more frequently

□ Yes: less frequently

□ No

17. has a measurement at your place of residence confirmed the presence of a physically source of low frequency sound?

□ Yes

□ No

□ Not done

18. does the perception of the humming sound represent a physiological or psychological stress for you?

□ Yes

□ No

19. what do you think is the cause of your perception of the humming sound?

□External sound source

□ Tinnitus

□ Spontaneous otoacoustic emissions

□ Sonstiges:________________________________________

20. how would you describe the quality of your perception of the humming sound?

□ Pure tone

□ Pulsating

□ Rising and falling

□ Constant

21 How old are you?

22. what gender are you?

□ Male

☐ Female

☐ Diverse
